# FOCUS: end-to-end preprocessing, alignment and resolution-matched integration of spatial multi-omics data

**DOI:** 10.64898/2026.08.04.742705

**Authors:** Lorenzo Venturelli, Jelle Jacobs, Alejandro Sifrim

## Abstract

**Summary:** Integrating spatial multi-omics data requires coordinated preprocessing, cross-modality alignment and feature registration across modalities that differ in file format, coordinate system and spatial resolution. No existing tool addresses this pipeline end-to-end from raw experimental files till aligned data object. We present FOCUS, an open-source Python package that takes raw data from spatial transcriptomics, mass spectrometry imaging, Raman spectroscopy imaging and brightfield or fluorescence microscopy through modality-specific preprocessing, interactive spatial alignment and resolution-matching registration to a unified MuData object, driven by a single configuration file. Its modular, registry-based architecture allows straightforward extension to additional modalities. FOCUS is accessible via a command-line interface, a browser-based GUI and a Python API.

**Availability and implementation:** FOCUS is implemented in Python 3.11, with a browser-based GUI built on a Vue.js 3 frontend served by a Flask backend. Source code, documentation and container recipes are available at https://github.com/sifrimlab/FOCUS; a versioned release is archived on Zenodo (10.5281/zenodo.21700038). Outputs use the AnnData and MuData formats and are directly compatible with the scverse ecosystem.

## 1. Introduction

A growing repertoire of spatial assays can profile a single tissue section across complementary molecular layers: gene expression by spot-based transcriptomics, lipid and metabolite distributions by mass spectrometry imaging, label-free chemical contrast by Raman spectroscopy imaging, and cellular morphology by brightfield or multiplexed-fluorescence microscopy, among others. The joint analysis of these layers can expose molecular relationships hidden to individual modalities, as demonstrated by recent spatially resolved studies combining transcriptomics and metabolomics on the same or serial tissue sections [1], [2]. Integrating these data requires all observations to be expressed on a common spatial index, which is technically demanding because the assays differ in file format, coordinate system, spatial resolution and preprocessing convention. When experimental constraints require serial sections rather than co-acquisition from the same physical section, inter-section biological and morphological variation introduces a further source of discrepancy.

The computational infrastructure for individual spatial modalities is well established. Scanpy [3], squidpy [4] and Giotto Suite [5] provide quality control, normalisation and analysis pipelines for spatial transcriptomics and multi-scale spatial omics data. Cardinal [6] and LipidQMap [7] covers preprocessing, peak alignment and tissue segmentation for mass spectrometry imaging. RamanSPy [8] provides spectral preprocessing routines for Raman imaging. For multiplexed histology, ASHLAR [9] performs tile stitching and cycle-to-cycle image registration, and VALIS [10] enables automated registration of whole-slide image series across staining protocols. Deep learning methods for automated spatial landmark detection, such as ELD [11], have been developed to identify correspondences across histology, transcriptomics and mass spectrometry imaging data. At the data level, SpatialData [12] and the muon framework [13] provide unified spatial data models and interfaces to downstream analysis tools, without themselves performing preprocessing, alignment or resolution matching. For aligning serial transcriptomics sections, PASTE [14], PASTE2 [15] and Stalign [16] infer spatial correspondences from expression similarity. SOmicsFusion [17] addresses co-registration between spatial metabolomics and biomedical imaging data, though without support for spot-based transcriptomics and currently distributed as standalone scripts. MAGPIE [2], a recently published Snakemake workflow, performs landmark-based co-registration of Visium [18] spatial transcriptomics and mass spectrometry imaging from the same or consecutive sections; it aggregates metabolite pixels onto the Visium spot grid and produces output compatible with established downstream analysis ecosystems. Both SOmicsFusion and MAGPIE accept preprocessed data as input, bypassing raw data conversion, and neither supports Raman spectroscopy imaging. No openly available tool addresses this pipeline end-to-end: from raw vendor files across dissimilar spatial technologies, through modality-specific preprocessing and cross-modality alignment, to a resolution-matched, analysis-ready integrated dataset.

Here we present FOCUS, an open-source Python framework for end-to-end preprocessing, spatial alignment and resolution-matching registration of multi-modal spatial omics data, from raw experimental files to a unified MuData object (Figure 1). FOCUS currently supports spatial transcriptomics, mass spectrometry imaging, Raman spectroscopy imaging and brightfield or fluorescence microscopy through a modular architecture designed to accommodate further modality types. It runs cross-platform and is operable through a command-line interface or a user-friendly browser-based graphical interface.

**Figure 1.**
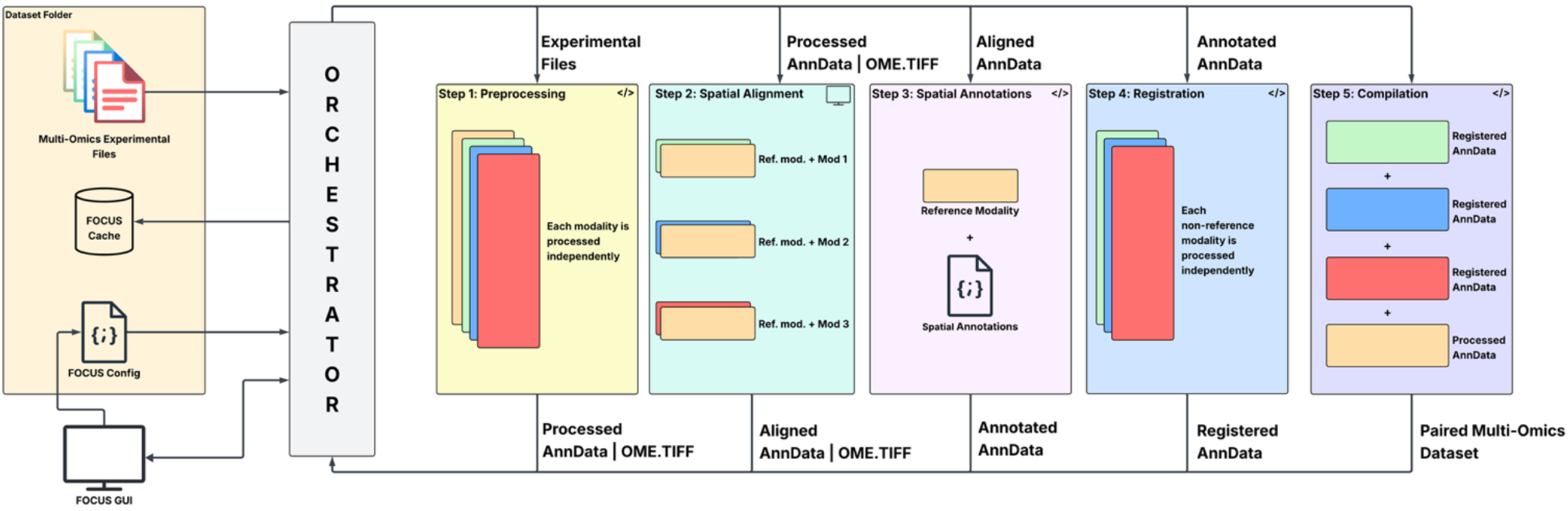
End-to-end schematic of the FOCUS pipeline. The dataset folder (left) supplies the raw multi-omics experimental files, an optional cache of intermediate outputs that enables resuming an interrupted run without recomputation, and a single JSON configuration file specifying the processing run (Section 2.1); these are read by the orchestrator regardless of whether the run is launched from the command line, the browser-based GUI or the Python API (Section 3). The orchestrator dispatches the dataset through five sequential stages, shown left to right: preprocessing, where each modality (colored boxes) is converted independently into a standardised AnnData or OME-TIFF representation (Section 2.2); spatial alignment, where each target modality (green, blue and red) is registered to the reference (orange) modality’s coordinate frame (Section 2.3); spatial annotation transfer, where GeoJSON region annotations are assigned to the reference observations (Section 2.3); registration, where each aligned modality is resampled or aggregated onto the reference observation grid (Section 2.4); and compilation, where the registered modalities are combined into the final paired multi-omics dataset (Section 2.5). Arrows above each stage label the data type passed forward as input to the next stage; arrows below label the corresponding output returned to the orchestrator before the next stage begins. Stacked boxes within a stage represent modalities processed independently of one another. The code icon (</>) marks stages executed automatically; the screen icon marks spatial alignment as the only stage requiring interactive input through the browser-based GUI (Section 4).

## 2. Methods

### 2.1 Multimodal data model and configuration

A FOCUS dataset is a collection of samples, each corresponding to one experimental unit such as a patient biopsy, a tissue replicate or an independent experiment. Each sample is profiled across one or more modalities; a modality denotes a single acquisition of the tissue by one technology. On disk, this hierarchy maps to a two-level directory tree: the dataset directory contains one subdirectory per sample, and each sample directory contains one folder per modality holding the raw experimental files. A single JSON configuration file per dataset encodes the full processing specification: which modalities to include, the preprocessing parameters for each, the designation of the reference modality, and the strategies for alignment and resolution matching. The file is human-readable and version-controllable, so the complete analytical protocol is self-contained, reproducible and distributable alongside the data. The reference modality defines the coordinate system and observation grid onto which all target modalities are mapped; it is typically the modality with the coarsest spatial resolution. Each coarse observation integrates signal over a spatial footprint that encompasses multiple fine-resolution locations, so mapping in the reverse direction would assign the same mixed measurement to several distinct positions, artificially inflating spatial resolution without recovering finer-grained information.

### 2.2 Modality-specific preprocessing

Each modality is preprocessed independently by routines tailored to its acquisition technology and converted into standardised intermediate representations: AnnData [1] .h5ad files for spot-based modalities and multi-resolution OME-TIFF pyramids for image-based ones. The AnnData format provides native compatibility with the spatial omics analysis ecosystem, and OME-TIFF is the community standard for large-scale bioimaging. Processing runs sample by sample, and per-sample outputs are merged into dataset-level files; both per-sample and merged files are persistent on disk so that an interrupted run can be resumed from any stage without recomputation. The modality-specific logic, covering operations such as background subtraction, illumination correction, spectral calibration, peak alignment and tissue segmentation, is described in Supplementary Methods.

### 2.3 Spatial alignment and annotation transfer

Spatial correspondence between modalities must be established explicitly, as instruments differ in coordinate origin, pixel scale and orientation. FOCUS addresses this through a browser-based alignment interface in which the reference layer is overlaid on each target layer and the user adjusts its position, orientation and shape until tissue structures visually agree. Critically, alignment requires no pre-defined landmark pairs: the user manipulates the overlay directly, making the approach applicable even when modalities lack clearly identifiable shared landmarks. The interface supports translation, rotation, scaling and homographic (per-corner) distortion, together accommodating the translation, rotation, scale and perspective differences that commonly arise between serial tissue sections. For modalities already sharing a coordinate frame with the reference, such as Visium spots recorded in the same H&E pixel coordinate system, the alignment step can be bypassed by declaring the modality as pre-aligned.

Spatial region annotations in GeoJSON format, such as pathologist-drawn tissue contours, are transferred to the reference observations by point-in-polygon assignment, with nested structures resolved by assigning each observation to its smallest containing polygon.

### 2.4 Spatial resolution matching

Once spatial correspondences are established, FOCUS maps each target modality onto the reference observation grid by computing a feature vector at every reference location. For spot-based targets, the target spots within each reference footprint are combined either as a normalized Gaussian-weighted average scaled to the reference spot size or, for subcellular-resolution assays, as a sum that accumulates rather than dilutes signal. The same Gaussian kernel is applied to Raman imaging targets, with the pixels of the stitched Raman image serving as the target spots. For microscopy image targets, a 224 × 224 pixel patch centred at each reference location is encoded by Prov-GigaPath [19], a pretrained pathology foundation model, into a 1,536-dimensional embedding vector. Reference locations without usable target content, whether due to an absence of nearby spots or a background-only image patch, are assigned zero vectors; this preserves row-to-row correspondence with the reference observation index and renders unmatched locations identifiable in downstream analysis. Prov-GigaPath weights are used as released by the original authors, without modification or fine-tuning. Spot interpolation runs on CPU. Image encoding runs on an NVIDIA GPU when CUDA is available and otherwise falls back to CPU; all remaining pipeline stages are CPU-only and run in parallel to remain tractable on standard workstations.

### 2.5 Integrated output

When the reference modality is spot-based, the aligned and resolution-matched modality files are assembled into a single MuData .h5mu object [13] in which every modality shares the reference observation index and spatial coordinates. This format is directly compatible with the muon [13] and scanpy [3] ecosystems for downstream analysis. When the reference modality is image-based, no MuData object is compiled: image targets are cropped to the aligned field of view and written as OME-TIFFs during image-to-image alignment, while spot-based targets are not integrated in this configuration and their outputs remain at the preprocessed stage.

## 3. Usage

FOCUS provides three execution interfaces, all of which consume the same configuration file and produce identical output. The command-line interface runs without a graphical environment and is suited to scripted and HPC workflows. Container recipes for Docker, Podman and Singularity/Apptainer support reproducible deployment on shared computing infrastructure without a local Python installation. The web GUI runs as a local web application and exposes the full pipeline: configuration building, validation, execution and per-stage progress monitoring. The alignment interface is only accessible through the GUI; command-line runs require that alignment has already been completed or that all target modalities are declared as pre-aligned.

All pipeline stages are additionally accessible through a Python API, which allows individual components to be called from notebooks or custom analysis scripts and embedded in larger computational workflows.

## 4. Limitations

Alignment in FOCUS is manual: visual correspondence must be established by the user through the browser-based GUI, which can be time-consuming for large cohorts and introduces operator variability. The geometric correction is restricted to a projective (homographic) warp, which cannot model local non-rigid deformations and may therefore reduce alignment accuracy for severely distorted tissue sections. Support for automated alignment, with approaches such as mutual-information optimization or learned feature matching, could be added as a complementary strategy in future releases without replacing the landmark-free manual approach.

At the dataset level, FOCUS requires every declared modality to be present in every sample; a run will fail if any sample is missing a declared modality file. At most one non-reference modality per run may be declared as pre-aligned. Both constraints limit applicability to complete and uniformly acquired datasets and may require manual curation of cohorts with uneven multi-modal coverage.

## Author contributions

LV: conceptualization and implementation of the framework, writing and editing of the manuscript.

JJ: discussions on conceptualization and testing of framework, editing the manuscript. AS: supervision of the work, editing of the manuscript

## Acknowledgements

LV is supported by the European Union’s Horizon 2020 research and innovation program under the Marie Skłodowska-Curie Actions Doctoral Network ‘PROSTAMET’, Grant Agreement No. 101120283 (HORIZON-MSCA-2022-DN-01).

JJ and AS are funded by the Leuven Future Fund (LISCO-BIOMED), Opening the Future Fund, KU Leuven ID-N (3E210655 COLUMBO).

## Conflict of interest

None declared.

## Supplementary Materials

### 1. Modality-specific preprocessing

Spatial transcriptomics samples are read as one AnnData [1] .h5ad file per sample, with raw counts in .*X* and coordinates in *obsm[‘spatial’]*, written as a compressed .h5ad; the reader is technology-agnostic. The pipeline flags mitochondrial genes by name, computes quality-control metrics including the mitochondrial fraction, and applies any per-spot count or gene filters the user requests. For the alignment interface it derives per-sample Leiden cluster labels on a spatially binned, internally normalized copy of the matrix, used only to color spots. Total-counts normalization, log1p transformation and mitochondrial-gene removal are all off by default, so .*X* retains raw counts and a raw layer is stored only when a normalization step would overwrite them. Samples are merged on disk by an outer join, filling absent genes with zeros; cross-sample gene filtering is optional and removes genes detected in too few samples, so by default every gene is kept, and the merged matrix is re-normalized only when normalization is enabled.

Mass spectrometry imaging samples are read as imzML/IBD pairs, optionally separated into positive- and negative-mode subdirectories for dual-polarity acquisitions, and written as a compressed .h5ad file per sample. The pipeline parses the imzML metadata; corrects the slight tilt of each acquisition’s laser grid relative to the discrete pixels’ coordinate axes, by fitting a simple linear regression model to the physical coordinates of its densest pixel column and rotating the coordinates back by the fitted angle. Additionally, for dual-polarity acquisitions, in which positive and negative modes are recorded separately and do not share a common coordinate frame, the two modes are registered to each other by fitting an affine transform between their physical coordinates and averaging the paired coordinates to one location per pixel. It then builds a consensus m/z grid across all spectra by adaptive sliding-window clustering at the mass tolerance, retaining clusters whose pooled weight reaches a set fraction of the largest weight identified across the samples. A small number of recalibration reference peaks per polarity is selected automatically from peak frequency and cross-sample coverage when none are supplied, per-spectrum m/z drift is corrected against them, and intensities are re-binned onto the consensus grid by inverse-distance-weighted interpolation. Foreground is separated from background on a composite spectral-complexity score inspired by the Cardinal package [20] combining Shannon entropy, peak count, total ion current and an optional lipid-database hit ratio. The score is than thresholded using a Gaussian mixture model with morphological cleanup for tissue samples or by an Otsu threshold with a percentile floor for microgrid samples. Intensity normalization is optional and defaults to none; the available modes are Total Ion Current (TIC), log-transform, Centered log-Ratio Normalization (CLR) and TIC Mean Scaled, with the CLR option centering each spectrum over its non-zero support and leaving structural zeros untouched.

Raman spectroscopy imaging samples are read as a single Leica .lif file per sample and written as a multi-resolution OME-TIFF. The pipeline parses the tile coordinates, pixel size, scan dimensions, spectral step count and pump-laser wavelength from the LIF XML and converts the spectral axis from emission wavelength to wavenumber using the pump wavelength. Illumination is corrected per channel with BaSiC, and background is segmented on a stitched single-component PCA mosaic through contrast-limited histogram equalization, Otsu thresholding, small-object removal and retention of the largest tissue contours. Each tile is then processed through a RamanSPy [8] pipeline of Whitaker-Hayes despiking, Savitzky-Golay denoising, IASLS baseline correction and min-max normalization, with zero-variance spectra flagged from the median absolute deviation of their forward differences. The corrected tiles are stitched with ASHLAR [9], using the highest-mean-intensity channel of the first cycle as the registration reference. BaSiC and ASHLAR run in two dedicated conda environments (FOCUS_BaSiCpy and FOCUS_ASHLAR) that are installation prerequisites for this modality.

Microscopy imaging samples are read as a single brightfield or fluorescence image per sample, in OME-TIFF, qpTIFF, plain TIFF or CZI format, and written as a multi-resolution OME-TIFF pyramid that preserves the input data type. The pipeline optionally enhances contrast, removes background by Otsu thresholding of a blurred, percentile-clipped image, cleans the resulting mask morphologically and crops to tissue. To stay tractable on large slides, the tissue mask is computed on a downsampled representation and broadcast back to full resolution. The stored pyramid keeps up to three channels, so inputs with more channels are truncated to their first three. This modality type is intended for imaging data that carry between one- and three-color channels.

### 2. Multimodal Alignment

Spatial correspondence between modalities is established in a browser-based alignment interface that loops over the samples each modality pair shares. The reference layer, whose observation grid all other modalities map onto, is drawn as a semi-transparent moving overlay above the fixed target layer, and the user adjusts it until tissue structures agree. The controls cover translation, rotation, isotropic and per-axis scaling, horizontal and vertical flips, and four-corner homographic distortion through draggable corner handles. No landmark correspondences are fitted: on confirmation the frontend returns a 3x3 transform, and the backend applies it to every reference observation and stores the result as obsm[‘{target_modality}_spatial’] on the reference object, expressing the reference observations in the target’s coordinate frame. For image targets the interface displays the lowest pyramid level and scales the confirmed coordinates back to full resolution; for image-to-image pairs it crops the reference to the aligned field of view and writes an OME-TIFF. A modality already in the reference frame, such as Visium spots recorded in their H&E pixel system, can instead be declared pre-aligned, which copies the coordinates without opening the interface; this requires a spot-based reference, accepts a target of any type, cannot be applied to the reference modality itself, and is limited to one non-reference modality per run (Supplementary Figure S1).

**Figure S1.**
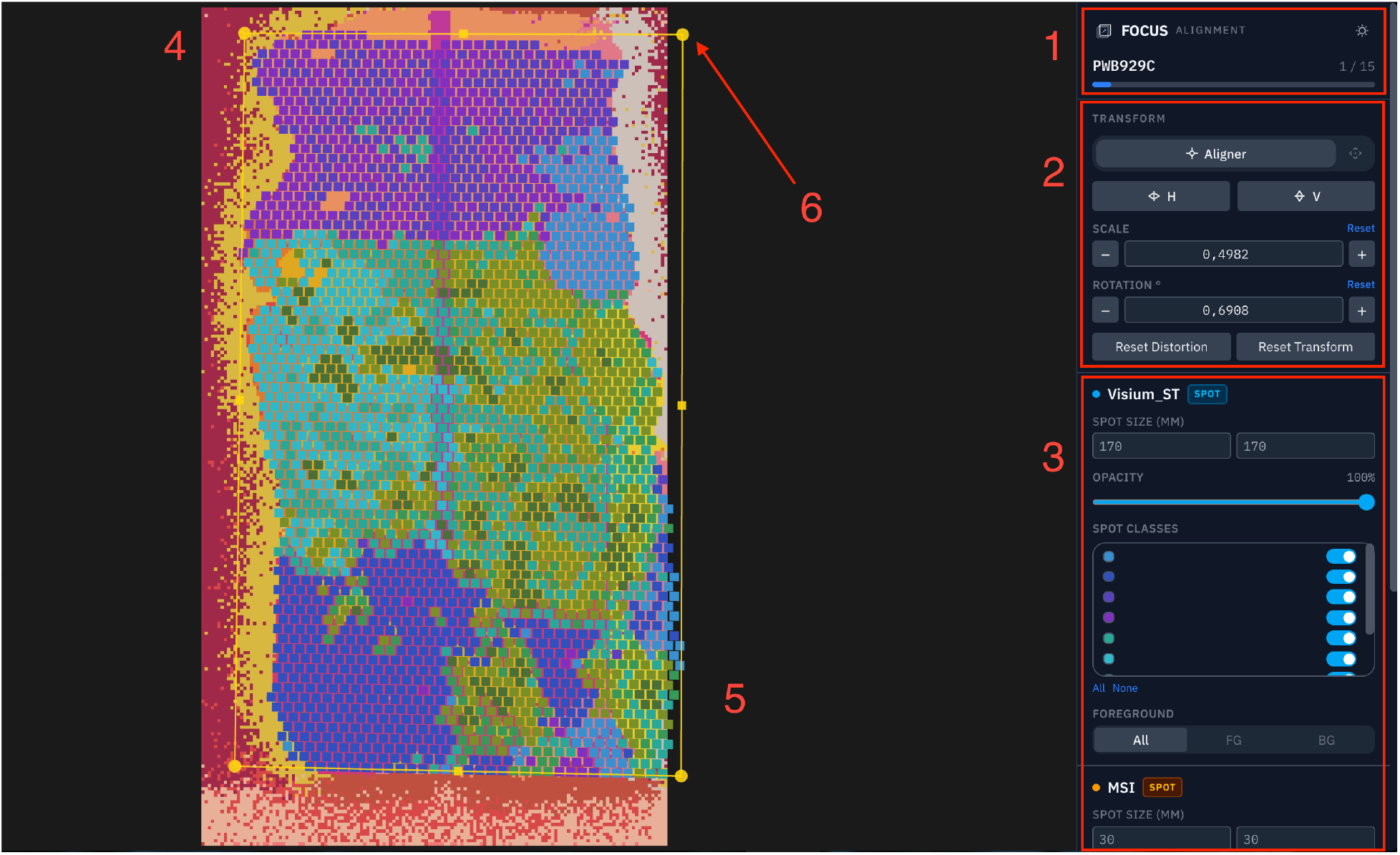
Annotated alignment interface. Screenshot of the browser-based alignment interface during registration of a mass spectrometry imaging (MSI) sample (target) to a 10x Visium spatial transcriptomics sample (reference). Both layers are rendered as point clouds, with each spot colored by its clustering label to highlight morphological structure. (1) Sample-pair indicator, showing the current sample index within the total number of samples shared by this modality pair. (2) Transform controls, including a toggle between alignment and camera-navigation modes, horizontal and vertical mirroring, scale and rotation adjustment, and transform reset buttons; these controls act on the reference layer. (3) Modality-specific display controls, including spot size, reference-layer opacity, per-cluster visibility, and toggling between the full point cloud and foreground-only or background-only points, available when background segmentation was performed for that modality during preprocessing. (4) Target modality (MSI), rendered as a fixed layer using a yellow-red color scale. (5) Reference modality (10x Visium spatial transcriptomics), rendered as a moving overlay using a blue-green color scale. (6) Bounding box of the reference layer: dragging the box translates the overlay, the scroll wheel scales it while in alignment mode and dragging a corner or edge handle applies rotation or perspective (homographic) distortion. No landmark correspondences are required for alignment.

### 3. Spatial Annotations

Region annotations are read from a GeoJSON FeatureCollection of Polygon and MultiPolygon features, with interior holes ignored. Each feature’s label is taken from its classification name, then its property name, then its feature identifier, whichever is available in this order. Polygons are tested against the reference observations by vectorised point-in-polygon queries on prepared geometries and assigned in descending order of area, so that where regions overlap the smallest containing polygon wins. The labels are written to obs[‘spatial_annotation’] on the reference modality and, after compilation, promoted to the top-level observation table of the integrated object.

### 4. Multimodal Registration

Registration maps each target modality onto the reference observation grid. The method is chosen per modality and constrained by the target’s type: Feature Extraction for *microscopy images*, Spot Interpolation or Spot Aggregation for the *spot-based modalities* (mass spectrometry imaging and spatial transcriptomics), and Raman Pixel Interpolation for *Raman*; a modality can also opt-out from the registration process. Reference locations left without usable target content receive zero vectors throughout, which keeps every registered modality in row-to-row correspondence with the reference index and marks unmatched locations for downstream filtering.

Feature extraction encodes microscopy morphology at each aligned reference location. A 224 x 224-pixel RGB patch centred on the location is passed through Prov-GigaPath [19], a pretrained pathology foundation model used as released, under ImageNet normalisation and in inference mode, producing a fixed-length (1,536-dimensional) embedding. Patches that are >99% background receive an all-zero embedding. Encoding runs on an NVIDIA GPU when CUDA is available, with the batch size set automatically from available memory, and falls back to a fixed batch on CPU. Due to the specificities of the pre-trained model, this registration step is currently only supported for haematoxylin and eosin (H&E) stained histological images.

Spot interpolation resamples a spot-based target onto the reference grid with a Gaussian kernel. For each reference location it queries a circular neighbourhood, keeps the target spots that fall within the reference footprint, and returns their Gaussian-weighted average, the kernel width scaled to the reference spot size. Locations with no target spot in the footprint receive a zero vector.

Spot aggregation shares the footprint geometry of spot interpolation but sums the target features within each footprint instead of averaging them and does not divide by the number of contributing spots. It is intended for subcellular-resolution assays such as Visium HD, where each native spot carries little signal and summation preserves it where averaging would dilute it. Visium HD is handled as a spatial transcriptomics modality, so the method applies to the spot-based modalities and needs no dedicated type.

Raman pixel interpolation applies the same Gaussian kernel with the pixels of the stitched Raman OME-TIFF as targets, each pixel contributing its spectral-intensity vector. The pixels are loaded only over a bounding box around the aligned reference locations, and locations that fall outside the image or capture no pixel receive a zero vector.

The integrated output depends on the reference type. When the reference is spot-based and at least one non-reference modality is registered, FOCUS assembles a single MuData [13] .h5mu object: the reference’s merged AnnData, annotated where annotation transfer was run, supplies the shared spatial coordinates, sample identifiers and spot size, and each registered modality is added only when its observation count and per-row sample order match the reference exactly, otherwise it is skipped. Reference observations that lack coverage in at least one target modality are dropped, so every retained observation is populated across all modalities, and each modality’s variable names are prefixed to prevent collisions on read-back. When the reference is image-based, no MuData object is compiled; cross-modality output in this configuration is the OME-TIFF cropped to the aligned field of view during image-to-image alignment, and spot-based targets remain at their preprocessed stage.

### 5. Extensibility through the modality and registration registries

FOCUS is extended through registries rather than by editing a central dispatcher. A new modality supplies a ModalityHandler (below) holding three callables: *create_samples* builds the per-sample processor objects from the dataset path and configuration; *create_dataset* assembles them into a dataset processor whose *process_dataset* method writes the per-sample and merged outputs; and *extract_settings* maps the modality’s configuration block to the keyword arguments those processors accept. A single *register_modality(modality_type, handler)* call adds the handler to the preprocessing registry, after which the orchestrator processes the new type without further changes to the pipeline. A new registration method follows the same pattern under *focus/registration/*: a class exposes *register_dataset(*…) and is dispatched by its *RegistrationType*. A few entries stay manual in either case. Adding a modality requires an entry in the file-extension, registration-compatibility, alignment-strategy-compatibility and display-name tables in *focus/constants*.*py*; adding a registration type requires a new *RegistrationType*, its compatibility and display-name entries, and a branch in the orchestrator’s registration dispatch. In both cases the browser configuration builder needs a matching option before the addition can be selected without hand-editing the configuration file. FOCUS validates configurations in code rather than against a schema file, so no separate schema needs updating.

@dataclass

class ModalityHandler:

“““Describes how to construct samples, dataset, and extract settings for a modality.”““

create_samples: Callable[…, list]

create_dataset: Callable[…, Any]

extract_settings: Callable[[dict], dict]

### 6. Functional comparison with related tools

Table S1 compares FOCUS with related tools across the capabilities that make up an end-to-end multimodal spatial-omics pipeline. FOCUS is the only tool in the comparison that spans the full path from raw experimental files, across dissimilar spatial technologies, through modality-specific preprocessing and cross-modality alignment, to a resolution-matched, analysis-ready integrated object. Tools built for a single stage or modality can be stronger within their scope: PASTE [14], PASTE2 [15] and STalign [16] align serial transcriptomics sections automatically, ASHLAR [9] and VALIS [10] register multiplexed and whole-slide microscopy, and Cardinal [6], [20] and RamanSPy [8] provide mature preprocessing for mass spectrometry and Raman imaging. Data frameworks such as SpatialData [12] and muon [13] supply unified containers but do not themselves preprocess, align or resolution-match. FOCUS partially relies on some of the mentioned tools to provide their capabilities inside a unified end-to-end workflow.

**Table S1:**
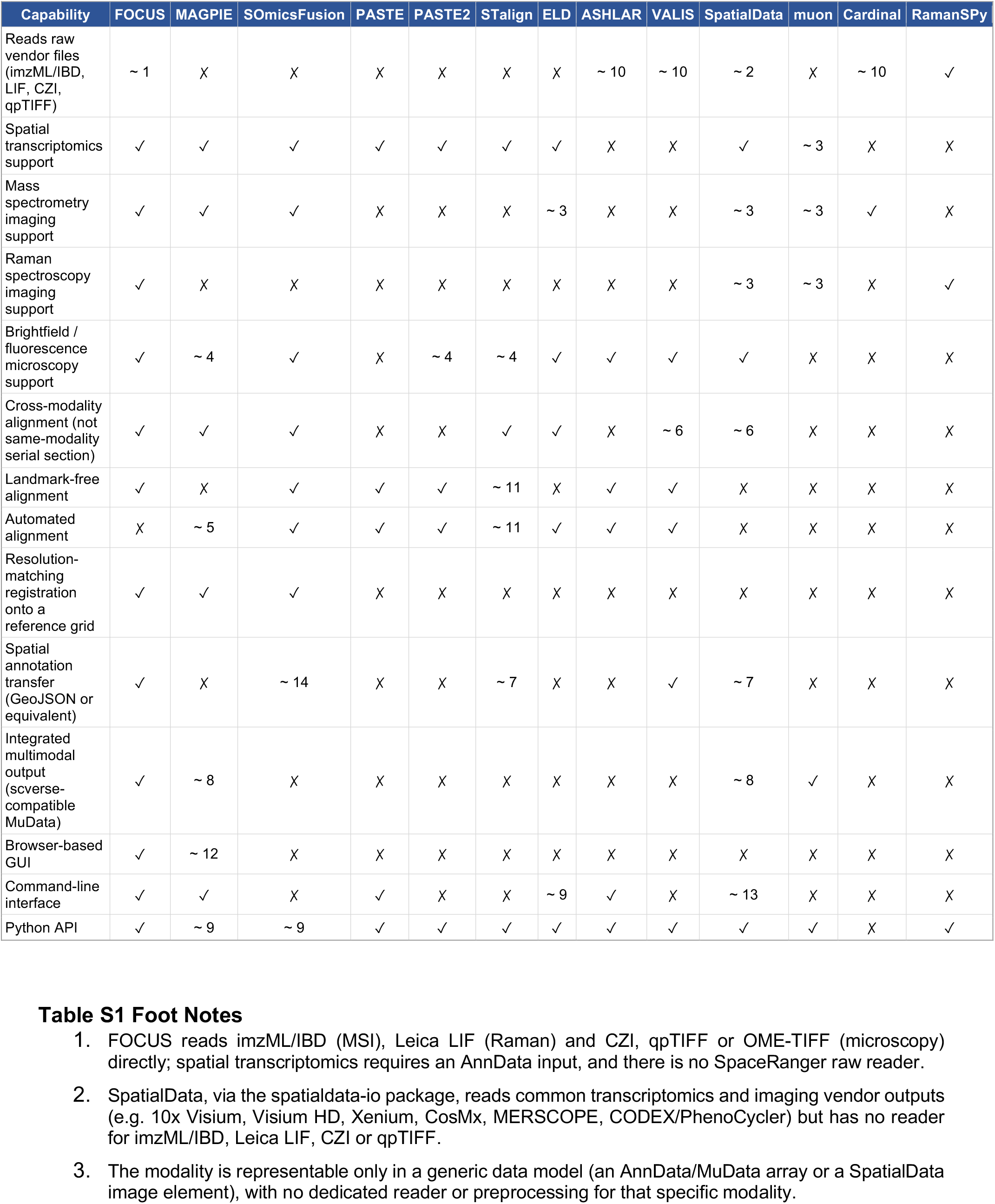
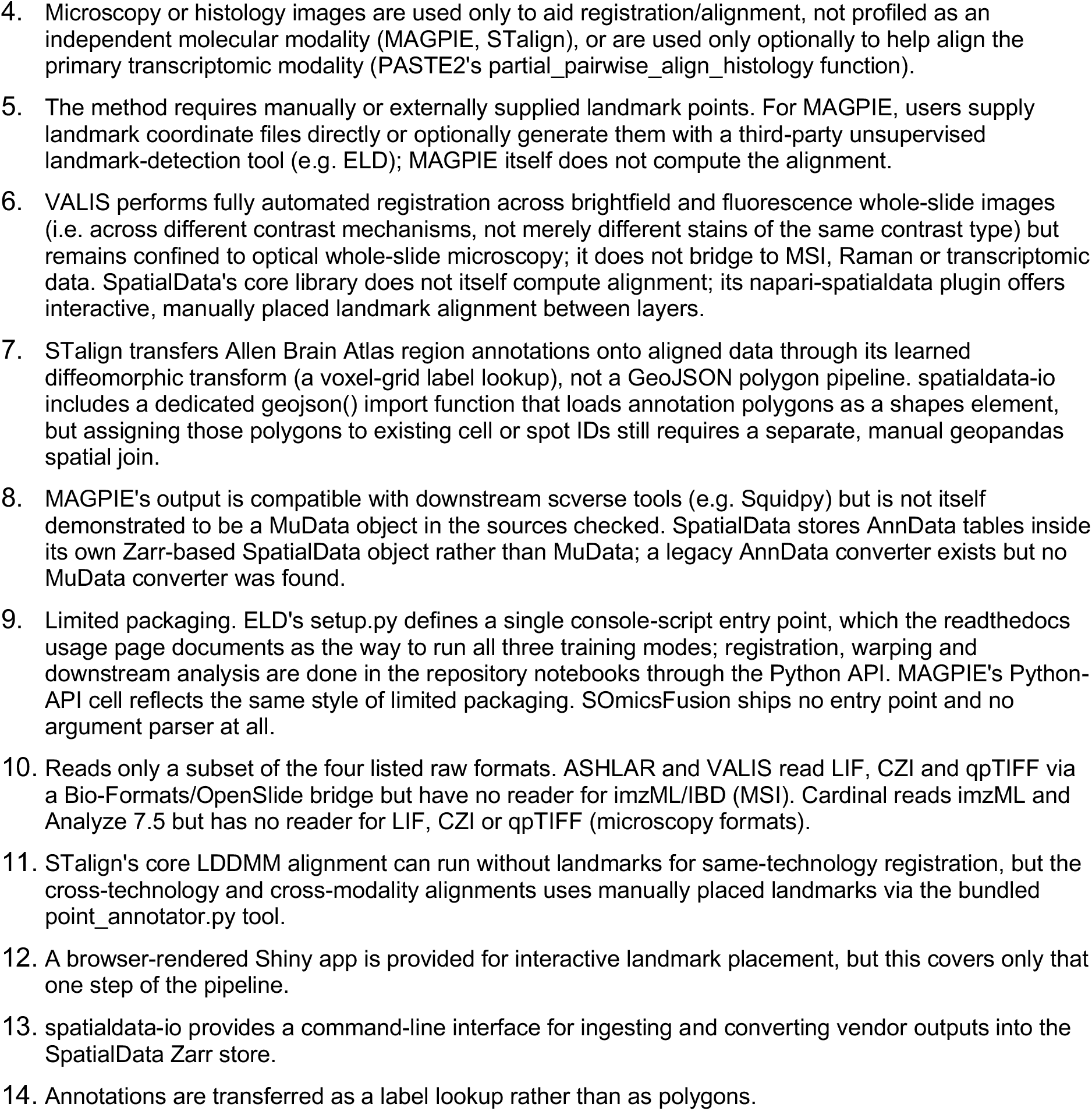
Functional comparison of FOCUS with related tools. Each functionality was assessed against the corresponding publication and the associated codebase. The ✓symbol denotes full support for the functionality, whereas ✗ denotes that the functionality is not supported. The ∼ symbol denotes partial or unverified support and is accompanied by a footnote providing further details

## Notes

### Competing Interest Statement

The authors have declared no competing interest.

## References

[1] M. Vicari et al., ‘Spatial multimodal analysis of transcriptomes and metabolomes in tissues’, Nat. Biotechnol., vol. 42, no. 7, pp. 1046–1050, Jul. 2024, doi: 10.1038/s41587-023-01937-y.

[2] E. C. Williams et al., ‘Spatially resolved integrative analysis of transcriptomic and metabolomic changes in tissue injury studies’, Nat. Commun., vol. 17, no. 1, p. 205, Jan. 2026, doi: 10.1038/s41467-025-68003-w.

[3] F. A. Wolf, P. Angerer, and F. J. Theis, ‘SCANPY: large-scale single-cell gene expression data analysis’, Genome Biol., vol. 19, no. 1, p. 15, Dec. 2018, doi: 10.1186/s13059-017-1382-0.

[4] G. Palla et al., ‘Squidpy: a scalable framework for spatial omics analysis’, Nat. Methods, vol. 19, no. 2, pp. 171–178, Feb. 2022, doi: 10.1038/s41592-021-01358-2.

[5] J. G. Chen et al., ‘Giotto Suite: a multiscale and technology-agnostic spatial multiomics analysis ecosystem’, Nat. Methods, vol. 22, no. 10, pp. 2052–2064, Oct. 2025, doi: 10.1038/s41592-025-02817-w.

[6] K. A. Bemis, M. C. Föll, D. Guo, S. S. Lakkimsetty, and O. Vitek, ‘Cardinal v.3: a versatile open-source software for mass spectrometry imaging analysis’, Nat. Methods, vol. 20, no. 12, pp. 1883–1886, Dec. 2023, doi: 10.1038/s41592-023-02070-z.

[7] J. Dehairs et al., ‘LipidQMap - An Open-Source Tool for Quantitative Mass Spectrometry Imaging of Lipids’, Oct. 15, 2025, Bioinformatics. doi: 10.1101/2025.10.15.682422.

[8] D. Georgiev, S. V. Pedersen, R. Xie, Á. Fernández-Galiana, M. M. Stevens, and M. Barahona, ‘RamanSPy: An Open-Source Python Package for Integrative Raman Spectroscopy Data Analysis’, Anal. Chem., vol. 96, no. 21, pp. 8492–8500, May 2024, doi: 10.1021/acs.analchem.4c00383.

[9] J. L. Muhlich, Y.-A. Chen, C. Yapp, D. Russell, S. Santagata, and P. K. Sorger, ‘Stitching and registering highly multiplexed whole-slide images of tissues and tumors using ASHLAR’, Bioinformatics, vol. 38, no. 19, pp. 4613–4621, Sep. 2022, doi: 10.1093/bioinformatics/btac544.

[10] C. D. Gatenbee et al., ‘Virtual alignment of pathology image series for multi-gigapixel whole slide images’, Nat. Commun., vol. 14, no. 1, p. 4502, Jul. 2023, doi: 10.1038/s41467-023-40218-9.

[11] M. Ekvall et al., ‘Spatial landmark detection and tissue registration with deep learning’, Nat. Methods, vol. 21, no. 4, pp. 673–679, Apr. 2024, doi: 10.1038/s41592-024-02199-5.

[12] L. Marconato et al., ‘SpatialData: an open and universal data framework for spatial omics’, Nat. Methods, vol. 22, no. 1, pp. 58–62, Jan. 2025, doi: 10.1038/s41592-024-02212-x.

[13] D. Bredikhin, I. Kats, and O. Stegle, ‘MUON: multimodal omics analysis framework’, Genome Biol., vol. 23, no. 1, p. 42, Dec. 2022, doi: 10.1186/s13059-021-02577-8.

[14] R. Zeira, M. Land, A. Strzalkowski, and B. J. Raphael, ‘Alignment and integration of spatial transcriptomics data’, Nat. Methods, vol. 19, no. 5, pp. 567–575, May 2022, doi: 10.1038/s41592-022-01459-6.

[15] X. Liu, R. Zeira, and B. J. Raphael, ‘Partial alignment of multislice spatially resolved transcriptomics data’, Genome Res., p. genome;gr.277670.123v1, Aug. 2023, doi: 10.1101/gr.277670.123.

[16] K. Clifton et al., ‘STalign: Alignment of spatial transcriptomics data using diffeomorphic metric mapping’, Nat. Commun., vol. 14, no. 1, p. 8123, Dec. 2023, doi: 10.1038/s41467-023-43915-7.

[17] A. Guo et al., ‘SOmicsFusion: Multimodal coregistration and fusion between spatial metabolomics and biomedical imaging’, Artif. Intell. Chem., vol. 2, no. 1, p. 100058, Jun. 2024, doi: 10.1016/j.aichem.2024.100058.

[18] P. L. Ståhl et al., ‘Visualization and analysis of gene expression in tissue sections by spatial transcriptomics’, Science, vol. 353, no. 6294, pp. 78–82, Jul. 2016, doi: 10.1126/science.aaf2403.

[19] H. Xu et al., ‘A whole-slide foundation model for digital pathology from real-world data’, Nature, vol. 630, no. 8015, pp. 181–188, Jun. 2024, doi: 10.1038/s41586-024-07441-w.

[20] K. D. Bemis et al., ‘Cardinal : an R package for statistical analysis of mass spectrometry-based imaging experiments’, Bioinformatics, vol. 31, no. 14, pp. 2418–2420, Jul. 2015, doi: 10.1093/bioinformatics/btv146.

